# Characterization of microsatellite markers for the duckweed *Spirodela polyrhiza* and *Lemna minor* tested on samples from Europe or the United States of America

**DOI:** 10.1101/2023.02.15.528655

**Authors:** Jae E. Kerstetter, Andrea L. Reid, Joshua T. Armstrong, Taylor A. Zallek, Trapper T. Hobble, Martin M. Turcotte

## Abstract

Microsatellite primers are a valuable tool to use for both observational and experimental studies in numerous taxa. Here, we develop 18 and 16 microsatellite markers for the widespread duckweeds *Lemna minor* and *Spirodela polyrhiza*, respectively. All 18 *L. minor* primers and 12 of the 16 *S. polyrhiza* primers amplified polymorphic loci when tested on samples from Europe or Western Pennsylvania, USA.

## Introduction

The globally distributed duckweed family (Lemnaceae) or subfamily (Lemnoideae) is composed of 36 species (Bog *et al*. 2020) of very small floating or submerged aquatic plants (Landolt, 1986; Sree *et al*. 2016). Duckweeds have a long history of scientific study given their highly specialized morphology, widespread distribution, high abundance, and production of the world’s smallest flowers (Jacobs, 1947; Hillman 1961; Landolt, 1986; Landolt, 1992). More recently, there has been an explosion in research interest given their potential applied uses including for agricultural feed (Cheng and Stomp, 2009), bioremediation (Gupta and Prakash, 2013; Ekperusi *et al*. 2019), and biofuel production (Cui and Cheng, 2015). Furthermore, their use as a model system to experimentally study numerous topics in ecology and evolutionary biology is quickly expanding (Laird and Barks, 2018). This growing basic and applied interest stems from their ability to reproduce clonally very quickly with population doubling times in as littles as 1.5 days (Ziegler *et al*. 2015). In addition, they are amenable to large scale manipulative experiments in both the lab and field mesocosms (Armitage and Jones, 2019; Hart *et al*. 2019; Tan *et al*. 2021; O’Brien *et al*. 2022), and have growing genomic data and tools (Wang *et al*. 2014; Ho *et al*. 2019; Xu *et al*. 2019; Cao *et al*. 2020) and characterization of their microbiome and herbivore communities (Acosta *et al*. 2020; Subramanian and Turcotte, 2020). Finally, duckweed express variation in numerous traits across species and among genotypes (clonal lineages) within species (Van Steveninck *et al*. 1992; Hart *et al*. 2019; Chen *et al*. 2020; Hitsman and Simons, 2020; Anneberg *et al*. 2023). Therefore, being able to identify genotypes may also be beneficial in many ecological studies to assess differences in traits among genotypes and to determine how these genotypes may respond to different environmental conditions.

Genetic markers, such as microsatellite markers, are important tools to study population genetics. Microsatellites, also known as simple sequence repeats ‘SSRs’, are tandem repeats two to 10 base pairs in length, that are flanked by conserved sequences and occur ubiquitously throughout eukaryotic genomes (Tautz and Renz, 1984). They are highly informative as locus-specific genetic markers due to their high abundance, high reproducibility, co-dominance, and polymorphic nature (Morgante and Olivieri, 1993; Powell *et al*. 1996). The length of the sequence repeats can be determined through PCR amplification using primers specific to their flanking regions; variation in PCR product length is a function of the number of repeated sequences. The high levels of polymorphisms observed in SSR markers (Tautz, 1989; Schlötterer and Tautz, 1992) and the relative ease of detection of these polymorphisms by PCR amplification has led to the wide applications of microsatellites as genetic markers (Vieira *et al*. 2016). Such within species markers have numerous applications including quantifying biogeographic distributions, population genetic structure, evolutionary history, and mating systems.

Moreover, a growing number of experimental evolution studies use such markers to track changes in genotypic composition of asexually reproducing populations over multiple generations (e.g., Turcotte *et al*. 2011; Agrawal *et al*. 2013; Hart *et al*. 2019) in large replicated experiments for which genotype-by-sequencing remains too costly. These cost savings are magnified when several loci can be genotyped in the same reaction (multiplexed; Markoulatos *et al*. 2002). Here, we report on the development of new microsatellite markers for two commonly studied and widespread duckweed species: the common duckweed *Lemna minor* (L. Schleid) and the greater duckweed *Spirodela polyrhiza* (L. Schleid)

With the growing interest in duckweed, microsatellite markers have been developed for a few duckweed species. Wani *et al*. (2014) developed nine polymorphic and 24 monomorphic haplotype cpDNA-based microsatellite primers for *L. minor*. Xu *et al*. (2018) developed 60 microsatellite primers for *Spirodela polyrhiza*, 19 of which were polymorphic within three populations of *S. polyrhiza* from China. Feng *et al*. (2017) developed three microsatellite primers for the identification of *S. polyrhiza* and *Landoltia punctata* haplotypes. More recently, Fu *et al*. (2020) developed 70 microsatellite primers within coding regions for *L. gibba*. It is important to continue developing and reporting new microsatellite markers as populations can differ in which markers function (e.g. due to null alleles) and are polymorphic (Chapuis and Estoup, 2007).

Here we report the successful development of 18 *L. minor* and 16 *S. polyrhiza* microsatellite markers. A small subset of these microsatellite primers were used to differentiate genotypes in our experimental studies on evolutionary-coexistence (Hart *et al*. 2019). In addition, we report genotyping results using these markers on individuals sampled in Europe and the United States of America (USA).

## Materials & Methods

### Sample Collection

Our objective when sampling was not to genetically characterize duckweed populations, but instead find genotypes that differ in ecologically relevant traits to use in various experiments. Thus, we genotyped few individuals from numerous bodies of water in various locations. Primers were developed at ETH Zurich (Europe) and the University of Pittsburgh (USA), and thus were tested on different collections of duckweeds. We collected duckweeds from numerous still bodies of water (e.g. ponds, lakes, wetlands) primarily in Switzerland and Western Pennsylvania (USA), however a few samples were also collected from the Netherlands and Germany. In addition, some European duckweed samples were obtained from the Landolt Duckweed Collection (formerly in Zurich, Switzerland) were included (see Supplemental Tables S1 and S2 for collection locations). Given the two-part development of the primers, some duckweed samples were only tested on the primers developed in that country (as noted in Tables S1 & S2).

Duckweeds mostly reproduce clonally via meristematic pockets from which clonal daughters emerge, creating clonal clusters of 1-8 individuals that eventually split into smaller clusters (Landolt, 1986). We sampled single duckweed clusters and established isofemale laboratory colonies from these clusters. We then sterilized each colony using sodium hypochlorite following a method adapted from Barks *et al*. (2018). From each colony, we put single individuals into individual sterile petri dishes (one individual per dish) containing sterile 0.5 strength Schenk and Hildebrandt growth medium (containing macro- and micronutrients as described by Schenk and Hildebrandt, 1972) supplemented with sucrose (6.7 g/L), yeast extract (0.067 g/L), and tryptone (0.34 g/L) for 24 hours to encourage algal and bacterial spore germination. Then each individual was exposed to one of an array of concentrations of sodium hypochlorite (0.3% or 0.5%) for varying amounts of time (3 or 6 minutes for *L. minor*, 4 or 7 minutes for *S. polyrhiza* respectively), then rinsed with autoclaved distilled water and allowed to grow (Barks *et al*. 2018). Sterile colonies were maintained in sterile 0.25 strength Schenk and Hildebrandt media (1972) without the additional supplements in room temperature laboratories or growth chambers under plant grow lights. These collections do not reproduce sexually under lab conditions.

### Microsatellite Marker Development

A total of 18 *L. minor* and 16 *S. polyrhiza* microsatellite markers were developed across Europe or the USA. We downloaded the whole genome shotgun sequence data for *S. polyrhiza* strain 7498 from the National Center for Biotechnology Information’s GenBank database (accession ATDW01000001.1) deposited by Wang *et al*. (2014). For *L. minor*, a draft genome (strain 8627) was downloaded from www.lemna.org on October 16th 2015 (genome draft lm8627.ASMv0.1) A recent study using Tubulin Based Polymorphism suggests that this lineage is in fact an interspecific hybrid of *L. japonica* and *L. turionifera* both closely related to *L. minor* (Braglia *et al*. 2021). The species identity for most samples on which we report below have been confirmed using morphology and/or barcoding (Fazekas *et al*. 2012; Barks *et al*. 2018). While some microsatellite markers are known to amplify across more than one duckweed species (Xu *et al*. 2018), we have not yet explicitly tested these markers against other species. Using MSATCOMMANDER (version 1.0.8, Faircloth 2008), we identified microsatellite loci using the default settings, except we avoided mononucleotide repeat motifs. We then selected loci that would produce products of different lengths, had different motif lengths, and were found on different contigs. The 5’ end of forward primers were labeled with one of several fluorescent dyes from various suppliers.

Primers developed at ETH Zurich were M13-tailed to reduce cost during development (Boutin-Ganache *et al*. 2001). This entailed adding the full or a partial M13 sequence of TGTAAAACGACGGCCAGT for the *S. polyrhiza* primers and GGAAACAGCTATGACCAT for *L. minor* primers to the 5’ end of the forward primer. The M13-labeled forward primers were used in combination with a M13 primer that had the same sequence but was fluorescently dye-labeled at its 5’ end. Some primers amplified less polymorphic loci or did not amplify as consistently as others. For these primers, we only have fragment lengths that include the M13 tail (see Tables 1 & 2), and we estimate that this lengthens the PCR product by 12-19 base pairs. For most primers however, following initial testing with M13, we ordered new labeled primers that did not include the M13 tail.

**Table 1:**
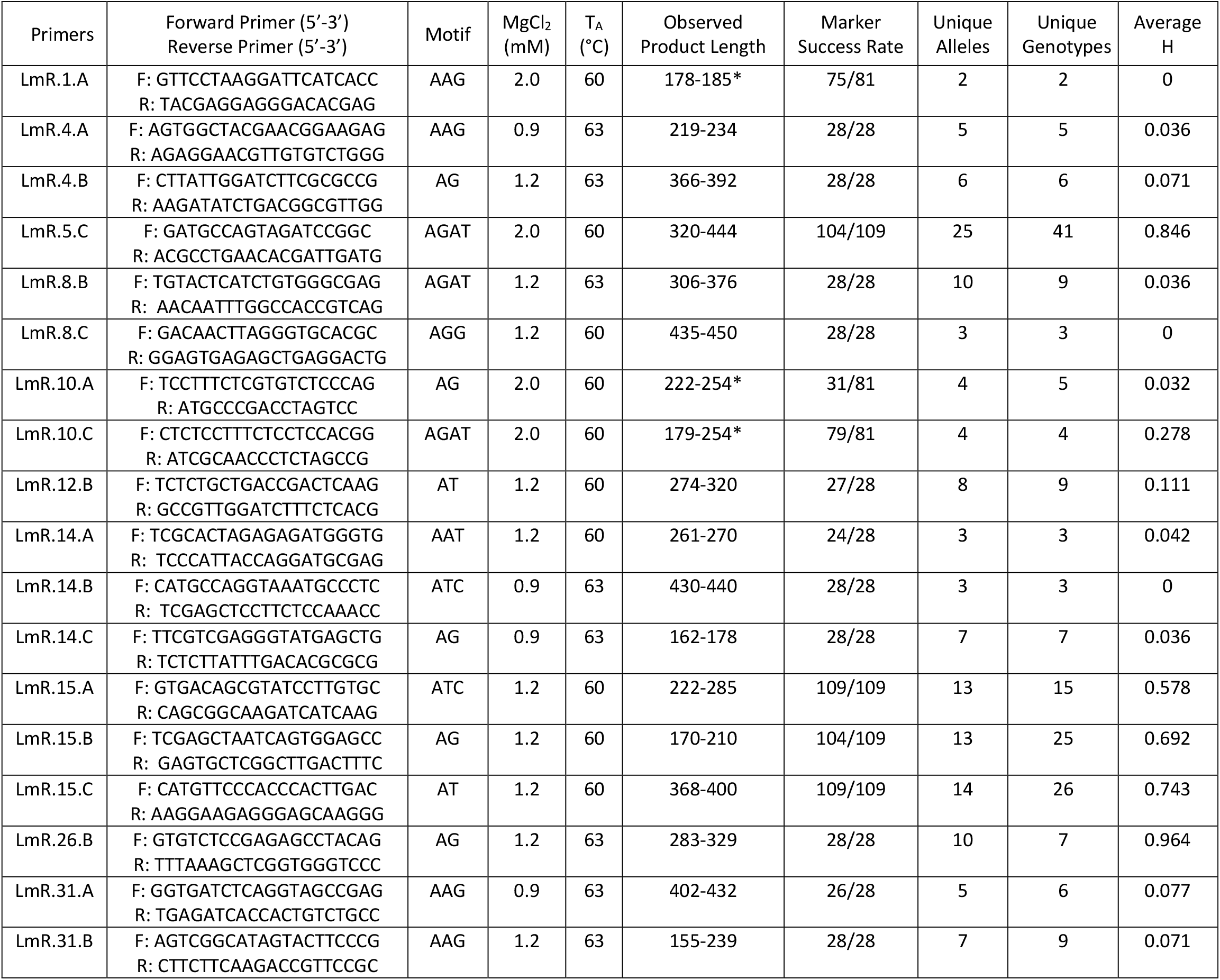
*Lemna minor* microsatellite markers and motifs including optimized MgCl2 concentrations and annealing temperatures (TA). In addition, we report marker success rate which is the number of samples successfully genotyped divided by those attempted, the number of unique alleles, and number of unique genotypes for each primer. Average heterozygosity (H) is the fraction of individuals that are heterozygotic for each primer. See Table S1 for specific allele values. Alleles lengths with an * denote that these lengths include the M13 tail sequence.

**Table 2:**
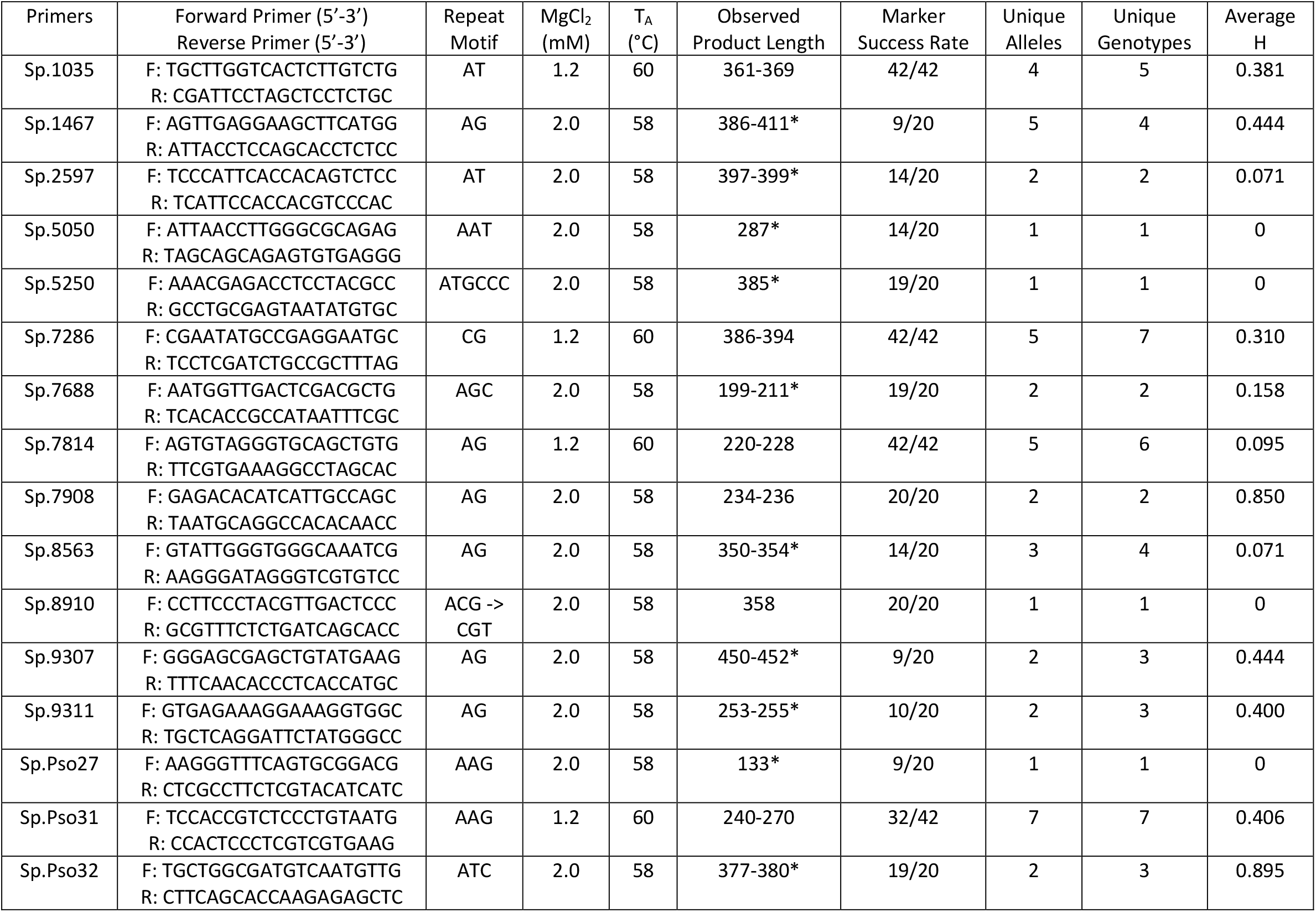
*Spirodela polyrhiza* microsatellite markers and sampling results as described in Table 1 with allele calls in Table S2.

At least 20 duckweed samples were tested using each primer. European duckweed samples were tested across 7 *L. minor* and 16 *S. polyrhiza* primers, and USA duckweed samples were tested across 15 *L. minor* and 4 *S. polyrhiza* primers (see Tables S1 and S2 for details).

### Microsatellite Amplification and Optimization

All duckweed collections were extracted and genotyped at least twice by first sampling 4 to 10 individuals from each monoclonal collection and lyophilizing them for 24 hours. We then extracted DNA using a modified CTAB-based method by Healey *et al*. (2014). Each microsatellite primer was first optimized for annealing temperatures and Magnesium chloride (MgCl2) concentration. The most commonly used PCR recipes and thermocycler conditions are presented here, and any primer specific differences are listed in Tables 1 and 2. For primers developed in Europe, the conditions were the following: PCR amplification was conducted in 15 μL volume reactions containing 3 μL of template DNA, 3μL of 5X Colorless GoTaq Flexi buffer (Promega, USA), 2.0 mM MgCl2, 0.2 mM dNTP mix, 0.05 μM of forward primer, 0.2 μM of reverse primer, 0.2 μM of M13 tagged with a fluorescent probe (e.g.: 5’ 6-FAM or 5’ HEX), and 1 unit of GoTaq G2 Flexi DNA Polymerase (Promega, USA). DNA concentrations was rarely quantified as amplification was often successful across a range of values (e.g. 2-40 ng/μL). Thermocycling conditions for both *S. polyrhiza* and *L. minor* from Europe that were M13 tagged were: initial denaturing at 94°C for 5 min, followed by 30 cycles of 1 min at 94°C, 1 min at 60°C, 1 min at 72°C, followed by eight M13 cycles consisting of 1 min at 94°C, 1 min at 53°C, 1 min at 72°C, followed by a final extension at 72°C for 10 minutes. For all primers developed in the USA, the conditions were the following: PCR amplification was conducted in 15 μL volume reactions containing 3 μL of template DNA, 3μL of 5X Colorless GoTaq Flexi buffer (Promega, USA), 1.2 mM MgCl2, 0.2 mM dNTP mix, 0.08 μg/μL of Bovine Serum Albumin (BSA), 0.2 μM of each forward and reverse primer, and 1 unit of GoTaq G2 Flexi DNA Polymerase (Promega, USA). Thermocycling conditions for *S. polyrhiza* were: initial denaturing at 94°C for 5 min, followed by 34 cycles of: 1 min of denaturing at 94°C, 1 min of annealing at 60°C, and 1 min of extension at 72°C, followed by a final extension at 72°C for 10 minutes. For *L. minor*, touchdown PCR was employed: with an initial denaturation of 94 °C for 5 min, followed by five cycles of denaturation (94 °C, 1 min), annealing (67 °C, 1 min; decreasing by 1 °C per cycle), and extension (72 °C, 1 min). Then 25 cycles of 1 min at 94°C, 1 min at 63°C, and 2 min at 72°C, followed by a final extension at 72°C for 15 minutes.

Fragment length analyses for all primers were conducted on ABI 3730 Genetic Analyzers (Applied Biosystems) at either the ETH Zurich Genetic Diversity Center (Switzerland), Keck DNA Sequencing Lab at Yale University (USA), or the University of Pittsburgh Genomics Research Core (USA), using either GeneScan™ 500 or 600 LIZ™ Dye Size Standards (Applied Biosystems). Allele calls were made using either Geneious (version 9.1.6, Keaser *et al*. 2012) or GeneMarker software (version 3.0.0, SoftGenetics, State College, Pennsylvania).

## Results and Discussion

We successfully developed 18 *L. minor* and 16 *S. polyrhiza* microsatellite primers (Tables 1 & 2) which were tested on samples of duckweeds from Europe or Western Pennsylvania (USA). Some markers were easier to utilize than others (Tables 1 & 2). All markers amplified in some samples; of these, all 18 *L. minor* primers and 12 of the 16 *S. polyrhiza* primers amplified polymorphic loci, having more than one allele. Moreover, these polymorphic loci differ in product length and can be used in multiplex reactions to increase efficiency and lower genotyping costs. We also found that some loci were much more polymorphic than others. For *L. minor*, these include loci amplified by primers LmR.5.C, LmR.8.B, LmR.15.A, LmR.15.B, LmR.15.C, and LmR.26.B, some of which showed high allele richness even when tested on only 28 samples (Table 1). For *S. polyrhiza* these included loci amplified by primers Sp.1467, Sp.7286, Sp.7814, and Sp.Pso31 (Table 2). Monomorphic loci may still be useful in different duckweed populations (Chapuis and Estoup, 2007). Many microsatellite loci also showed heterogeneity (Tables S1 and S2) which helps make the primers more informative to distinguish genotypes. We note that some primers developed in one continent were not tested on samples from the other continent (see footnotes in Tables S1 &S2); we suspect these primers will work across continents given patterns observed in the others, but this remains to be tested.

Comparing between species, we see that although we tested less samples of *S. polyrhiza*, it still has much lower allelic and genotypic richness across most primers. This is consistent with our own recent large-scale sampling (Hobble *et al. In prep)* as well as other studies using different genotyping methods, that similarly found low genetic diversity in *S. polyrhiza* (Bog *et al*. 2015; Xu *et al*. 2015; Feng *et al*. 2017). It has been hypothesized that this low genetic variation in *S. polyrhiza* is due to its low mutation rate (Xu *et al*. 2019). In addition, primers differed greatly in average observed heterozygosity, but species had similar mean heterozygosities (0.256 for *L. minor* and 0.283 for *S. polyrhiza*). Given that our sampling was designed to find unique genotypes (shallow and widespread) and not characterize populations, we limit our discussion or quantification of population genetic indices. The primers we developed can help researchers address various ecological and evolutionary questions as well as better identify and catalogue genotypes for applied activities.

## Supplemental Table Captions

**Table S1:** *Lemna minor* sample collection sites and allele lengths.

**Table S2:** *Spirodela polyrhiza* sample collection sites and allele lengths.

## Acknowledgements

We thank Walter Lämmler for sharing duckweed lineages from the Landolt Duckweed Collection. We are grateful to the former Levine Plant Ecology Group at ETH Zurich and the Turcotte Lab for their assistance in maintaining collections. We thank ETH Zurich Genetic Diversity Center and Mary Janecka for help troubleshooting primer development. M.M.T. was supported by the ETH Zurich Center for Adaptation to Changing Environments and now by an NSF grant DEB-1935410.

## Notes

### Competing Interest Statement

The authors have declared no competing interest.

